# Growth impacts in a changing ocean: insights from two coral reef fishes in an extreme environment

**DOI:** 10.1101/2020.06.26.174029

**Authors:** Daniele D’Agostino, John A. Burt, Veronica Santinelli, Grace O. Vaughan, Ashley Fowler, Tom Reader, Brett M. Taylor, Andrew S. Hoey, Georgenes H. Cavalcante, Andrew G. Bauman, David A. Feary

**Affiliations:** School of Life Sciences, University of Nottingham, Nottingham NG7 2RD, UK; Center for Genomics and Systems Biology, New York University Abu Dhabi, PO Box 129188, Abu Dhabi, United Arab Emirates; Department of Earth and Marine Sciences, University of Palermo, Via Archirafi 18, 90123 Palermo, Italy; New South Wales Department of Primary Industries, Sydney Institute of Marine Science, Mosman, NSW, 2088, Australia; School of Life Sciences, University of Technology, Sydney, Broadway, NSW, 2007, Australia; Australian Institute of Marine Science, Crawley, Western Australia 6009, Australia; ARC Centre of Excellence for Coral Reef Studies, James Cook University, Townsville, 4811, Australia; Instituto de Ciências Atmosféricas, Universidade Federal de Alagoas, Maceió, Alagoas, 57072-970, Brazil; Department of Biology, Chemistry and Environmental Sciences, American University of Sharjah, PO Box 26666, Sharjah, United Arab Emirates; Experimental Marine Ecology Laboratory, National University of Singapore, 21 Lower Kent Ridge Road, 119077, Singapore; MRAG Ltd, 18 Queen Street, London W1J 5PN, UK

**Keywords:** ocean warming, TSR, coping mechanism, plasticity, stress, physiology, oxygen limitation

## Abstract

Determining the life history consequences for fishes living in extreme and variable environments will be vital in predicting the likely impacts of ongoing climate change on reef fish demography. Here, we compare size-at-age and maximum body size of two common reef fish species (*Lutjanus ehrenbergii* and *Pomacanthus maculosus*) between the environmentally extreme Arabian/Persian Gulf (‘Arabian Gulf’) and adjacent comparably benign Oman Sea. Additionally, we use otolith increment width profiles to investigate the influence of temperature, salinity and productivity on the individual growth rates. Individuals of both species showed smaller size-at-age and lower maximum size in the Arabian Gulf compared to conspecifics in the less extreme and less variable environment of the Oman Sea, suggesting a life-history trade-off between size and metabolic demands. Salinity was the best environmental predictor of interannual growth across species and regions, with low growth corresponding to more saline conditions. However, salinity had a weaker negative effect on interannual growth of fishes in the Arabian Gulf than in the Oman Sea, indicating Arabian Gulf populations may be better able to acclimate to changing environmental conditions. Temperature had a weak positive effect on the interannual growth of fishes in the Arabian Gulf, suggesting that these populations may still be living within their thermal windows. Our results highlight the potential importance of osmoregulatory cost in impacting growth, and the need to consider the effect of multiple stressors when investigating the consequences of future climate change on fish demography.

## Introduction

Anthropogenic global climate change is having significant biological impacts on individuals, species and ecosystems (e.g. Parmesan and Yohe 2003; Bellard 2012; Hughes et al. 2018; Pratchett et al. 2018; França et al. 2020). Reductions in body size, coupled with geographic shifts in species distribution (Walther et al. 2002; Parmesan and Yohe 2003; Feary et al. 2014) and changes in phenology (Walther et al. 2002; Stenseth 2002; Visser and Both 2005; Taylor 2008), are recognised as common responses to global warming in ectotherms (Daufresne et al. 2009; Gardner et al. 2011; Ohlberger 2013). Such reduction in body size is consistent with the temperature-size-rule (TSR) (i.e. body size decrease at higher temperature) (Atkinson 1994; Kingsolver and Huey 2008; Ohlberger 2013; Horne et al. 2017; Huss et al. 2019) and is particularly evident in aquatic environments (Forster et al. 2012; Horne et al. 2015), where fish and other aquatic organisms’ average body size has already declined by 5–20% over the last two decades (Baudron et al. 2014; Audzijonyte et al. 2016; van Rijn et al. 2017). Experimental temperature-size responses, models and meta-analyses suggests that body size will further reduce by 3–5% per degree of warming in aquatic arthropods, while in fishes it may decline by 14–24% by 2050 under a higher green-house gases emission scenario (A2 scenario, IPCC 2007; Cheung et al. 2013; Pauly and Cheung 2018a).

Decreasing fish sizes will impact fecundity and fisheries productivity (Baudron et al. 2014; Barneche et al. 2018), prey-predator interactions (Barnes et al. 2010), and overall ecosystem functioning (Bellwood et al. 2012; Cheung et al. 2013). Understanding what drives reductions in fishes body size will be essential for predicting how populations are impacted by projected climate change. In this regard, despite the ubiquity of the TSR in explaining size reduction in a range of taxa, the underlying mechanisms of body size reduction with increasing temperature are still debated (Audzijonyte et al. 2019).

Projected increases in mean temperature and temperature variability due to climate change (i.e. 4.0 °C increase by 2100, IPCC 2014; Hoegh-Guldberg 2018) are likely to be particularly challenging for marine fish due to the greater energetic demands for routine metabolic activities (Gillooly et al. 2001; Pörtner and Knust 2007; Neuheimer et al. 2011) with, ultimately, less energy available for somatic growth. Similarly, projected intensification of the global water cycle (4–8% per degree Celsius of increase of surface air temperature change) and consequent increase of water salinity in already dry regions (Durack et al. 2012; Skliris et al. 2014; Zika et al. 2018), will likely increase the osmoregulation costs for many marine species (Boeuf and Payan 2001; Ern et al. 2014), especially those living in hypersaline semi-enclosed seas.

One approach to understanding how marine fish demography may be impacted by future environmental variance and extremes in temperature and salinity is to study contemporary communities that exist within naturally extreme and variable environments. Here, we compare the effect of water temperature, salinity and primary productivity on the size-at-age and growth rate of coral reef fishes between the environmentally extreme southern Arabian/Persian Gulf (hereafter ‘Arabian Gulf’) and comparatively more benign adjacent Oman Sea (Fig. 1). The Arabian Gulf has the largest thermal range (> 20 °C) and highest maximum sea surface temperature (SST) experienced by extant coral reef fishes (winter: < 15 °C, summer: > 35 °C), with fishes enduring several months of conditions considered lethal to reef fishes in other parts of the world (Riegl and Purkis 2012; Rummer et al. 2014; Vaughan et al. 2019). In addition, present summer SSTs in the Gulf are comparable to those expected for tropical oceans by 2100 (Riegl and Purkis 2012; Hoegh-Guldberg 2018), while winter temperatures can be so low as to induce cold water coral bleaching (Shinn 1976; Coles and Fadlallah 1991). Additionally, due to restricted water exchange through the narrow Strait of Hormuz, as well as limited freshwater input (Reynolds 2002) and high evaporation (Reynolds 1993), the Arabian Gulf’s waters are characterized by hypersaline conditions (annual mean salinity 42 psu) that are the highest reported for coral reefs (Sheppard 1993, Bauman 2013, Vaughan et al. 2019). In contrast, the Oman Sea is subject to less extreme and variable temperature and salinity, but a higher baseline and higher interannual variation of chlorophyll-a concentration (used as a proxy for primary productivity) (Coles 1997, Coles 2003).

**Fig. 1.**
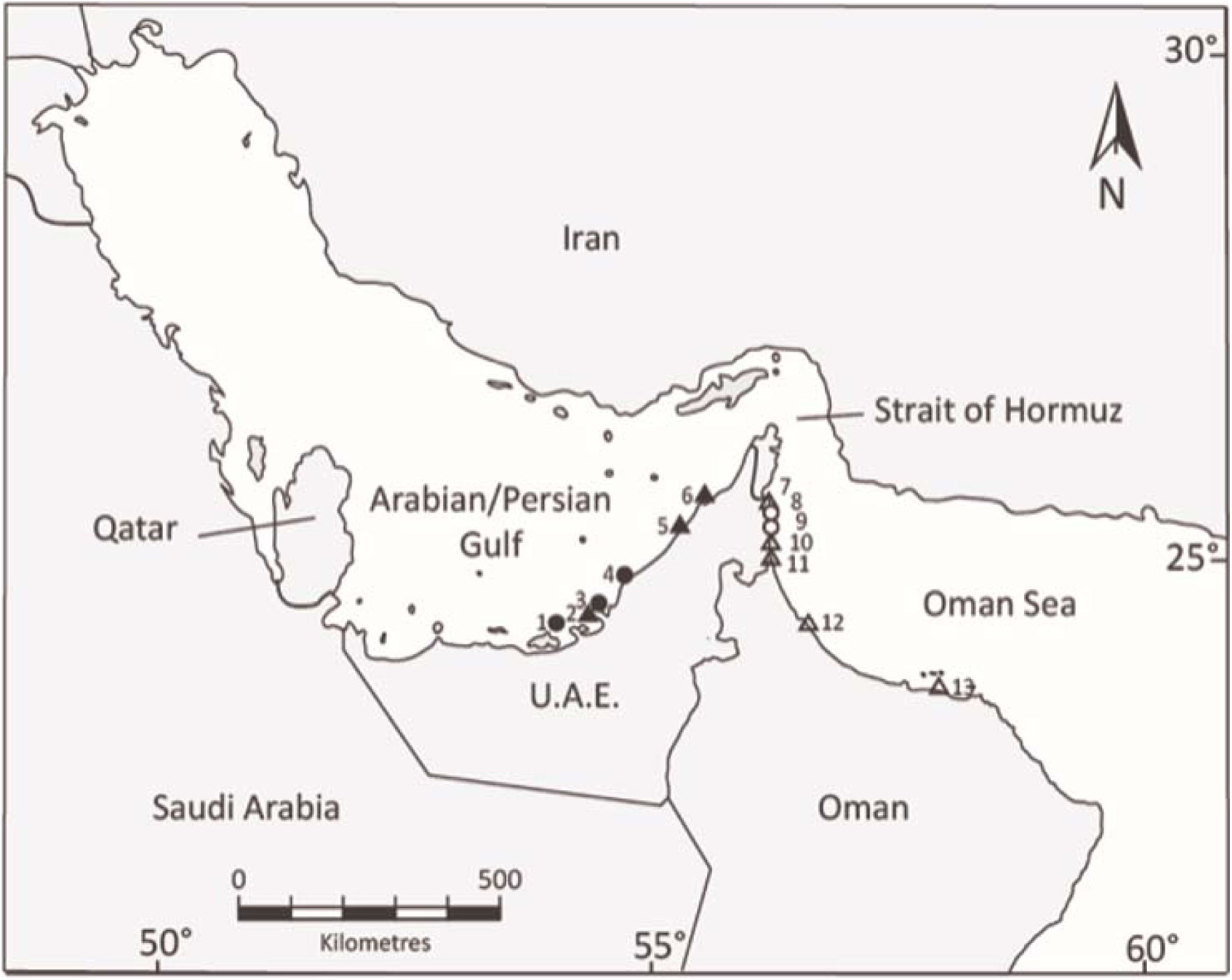
Map of the study regions showing collection locations in the southern Arabian/Persian Gulf and northern Oman Sea (circles: field collection–reefs, triangles: local fish landing sites; refer to Table S2 for location names, coordinates and sampling method).

Given differences in environment conditions between the Arabian Gulf and the Oman Sea, we hypothesise that, in line with the TSR, (i) fishes in the Arabian Gulf would exhibit lower growth rates, as well as smaller size-at-age and maximum body size than conspecifics from the less extreme and less variable environment of the Oman Sea; and (ii) that predicted differences in growth would be driven interannually by local environmental conditions. We investigated the role of temperature, salinity and productivity in structuring fish demography between regions by using an otolith sclerochronology approach and mixed-effect modelling, and predicted that low growth rate and small body size would be the results of extreme temperature ranges in the Arabian Gulf, exacerbated by high salinity and lower primary productivity.

## Materials and methods

To determine whether fish growth is reduced in the Arabian Gulf compared to the Oman Sea, and whether extreme environmental conditions are the drivers of somatic growth differences, we analysed the otolith structure of the blackspot snapper (*Lutjanus ehrenbergii*, F. Lutjanidae [Peters, 1869]) and the yellow-bar angelfish (*Pomacanthus maculosus*, F. Pomacanthidae [Forsskål, 1775]). Both *L. ehrenbergii* and *P. maculosus* are coral-associated species commonly found on nearshore patch and fringing reefs throughout both regions (Feary et al. 2010; Burt et al. 2011) and are known to have highly determinate growth patterns (Grandcourt et al. 2010; 2011). Furthermore, the two species were chosen because they are phylogenetically and ecologically disparate: *L. ehrenbergii* is a generalist carnivore, feeding on benthic invertebrates associated with turfing algae (i.e. amphipods, isopods) and small fish (Randall 1995; D’Agostino et al. 2019), while *P. maculosus* is a facultative spongivore/corallivore (Shraim et al. 2017).

### 2.2 Sample collection of focal species for life history comparison

Sample collection encompassed six sites within the Arabian Gulf (three reefs and four local fish landing sites, spanning from Dhabiya to Umm Al Quwain) and seven sites within the Oman Sea (two reefs and five local fish landing sites, spanning from Dibba to As Seeb) (Fig. 1, Table S2). As both species are relatively site attached and the nearest sites between the two regions were separated by more than 300 km (Grizzle et al. 2015), negligible movement between regions was predicted (Buchanan et al. 2019). Individuals collected *in situ* (i.e. from the reefs) were collected using the fish anaesthetic clove oil (juveniles) or spear guns (sub-adults and adults) and euthanized using an ice slurry. Upon collection, all individuals were measured (standard length [SL], nearest mm), weighed (total weight, nearest g) and sagittal otoliths removed, cleaned in ethanol and stored dry.

### 2.3 Age determination

To determine age, 1012 and 210 otoliths were sectioned from *L. ehrenbergii* and *P. maculosus* specimens, respectively (Table 1). Each otolith was ground to the nucleus to produce a thin transverse section and mounted on a microscope slide using established procedures (i.e. Taylor and McIlwain 2010). Otoliths were examined under transmitted light with a low power microscope and individual ages estimated by counting the number of annual increments, or annuli, along the dorsal antisulcus axis of each otolith (Fig. 2). Previous studies have verified the deposition of annual increments for both species (Grandcourt et al. 2010; 2011). Blind reads (for size and collection location) of annual increments were performed on three separate occasions for each specimen. Final age was determined when two or more counts agreed. If agreement was not achieved after three counts, the sample was excluded from the analysis. For individuals < 1 year, otolith sections were further ground with lapping film, polished by hand with 0.3 µm alumina powder and viewed through a compound microscope. For these individuals age was then estimated by counting the number of daily increments, with three blind reads performed for each individual. Final age for juvenile fishes was taken as the median of three counts when all counts were within 10% of the median. Samples with counts > 10% of the median were excluded from the analysis.

**Table 1.** Details of *Lutjanus ehrenbergii* and *Pomacanthus maculosus* otolith samples used to investigate differences in growth between the Arabian Gulf and Oman. Site and species-specific growth comparison and best-fit re-parameterised von Bertalanffy growth-function

| Species | Area | *n* | Size<br>range<br>(mm) | Age<br>(days □<br>years) | Age<sub>max</sub> (SE) | L<sub>max</sub> (SE) | L<sub>1</sub> | L<sub>3</sub> | L<sub>6</sub> | - $\lambda$ | $\sigma$ |
| --- | --- | --- | --- | --- | --- | --- | --- | --- | --- | --- | --- |
| *Lutjanus*<br>*ehrenbergii* | Arabian<br>Gulf | 873 | 33 □ 224 | 38 □ 15 | 8.58 (0.41) | 199.52 (1.91) | 132.54 | 186.31 | 189.45 | 4071.80 | 25.53 |
| | Oman | 139 | 18 □ 282 | 27 □ 16 | 11 (0.53) | 246.21 (3.71) | 144.72 | 216.52 | 223.12 | 695.66 | 34.81 |
| *Pomacanthus*<br>*maculosus* | Arabian<br>Gulf | 168 | 9 □ 262 | 9 □ 31 | 14.62 (1.14) | 197.48 (4.94) | 95.12 | 160.65 | 170.48 | 841.03 | 36.12 |
| | Oman | 42 | 40 □ 264 | 81 □ 22 | 8.55 (1.93) | 227.11 (6.55) | 107.34 | 177.83 | 188.34 | 204.78 | 28.65 |

parameters (rVBGF) are shown.
*n*, sample size; L<sub>1</sub>, mean size at age one; L<sub>3</sub>, age three; L<sub>6</sub>, age six; - $\lambda$ , negative log-likelihood; $\sigma$ , standard deviation. All sizes are in mm SL, Ages are in years unless indicated otherwise.

**Fig. 2.**
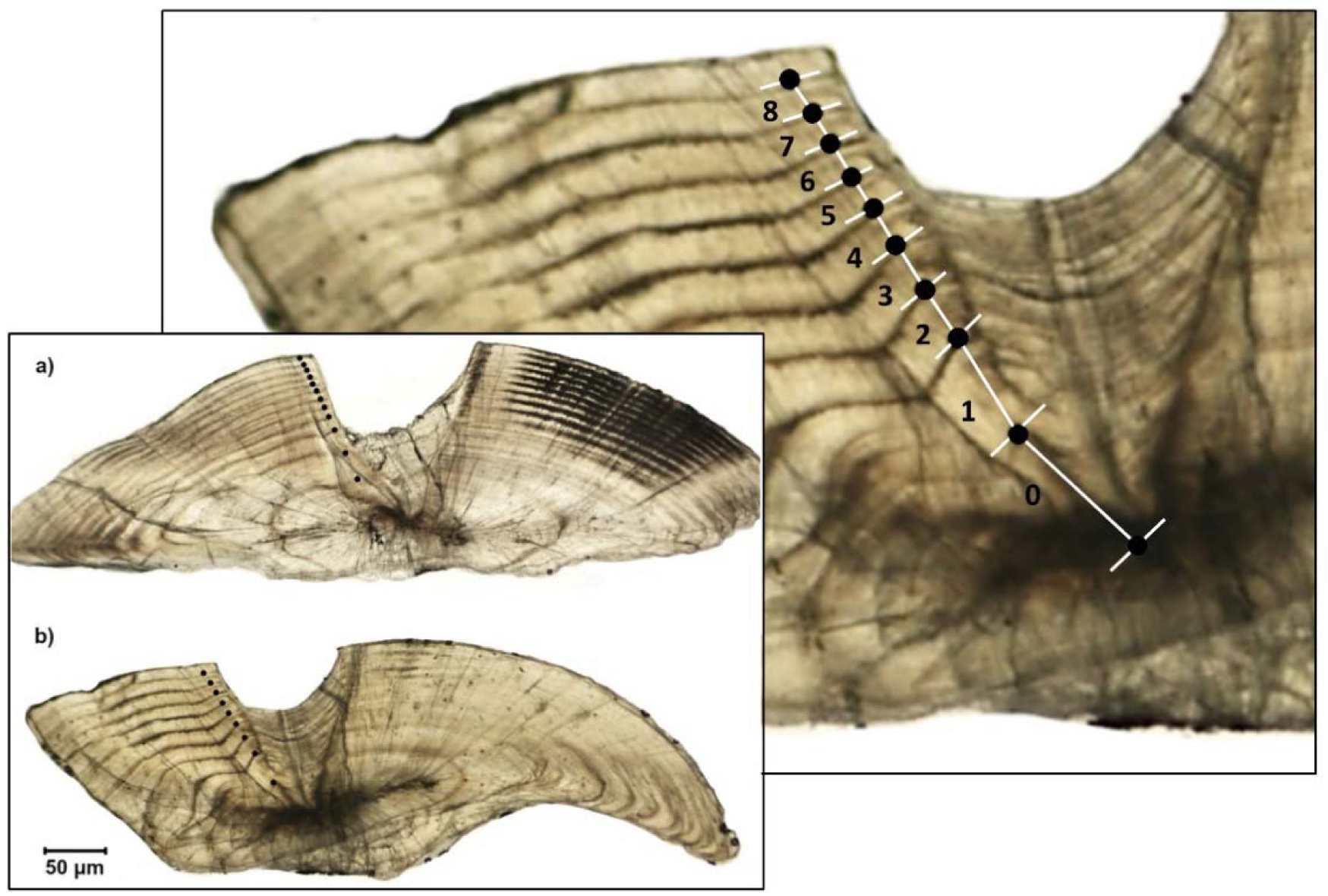
Annual increment widths follow measurements along transverse sections of otoliths in **(a)** *Lutjanus ehrenbergii*, and **(b)** *Pomacanthus maculosus*. Photographs were taken under a high-power microscope with transmitted light. Inset shows the annotated transverse section of a fish estimated as 8 years old.

### 2.4 Population-level growth, body size and life span

To model and compare the population growth rate of *P. maculosus* and *L. ehrenbergii* between the Arabian Gulf and the Oman Sea, the re-parameterised version of the von Bertalanffy growth function (rVBGF, Francis 1988) was used (Visconti et al 2018). The rVBGF describes growth based on modelled body size of individuals at three ages, L_*τ*_, L_*ω*_ and L_*μ*_, allowing a direct comparison of mean size-at-age data between populations. Age *τ* represents fast, early growth (i.e. ascending part of the growth trajectory), *μ* represents the asymptote of growth (i.e. when growth reaches a plateau), while *ω* represents onset of a reduction in growth rate (Trip et al. 2008, 2014). To model species growth based on the trajectory of the region-specific growth curves, ages 1 and 6 years (based on annuli counts) were taken as L_*τ*_(hereafter ‘L_1_’) and L_*μ*_(hereafter ‘L_6_’), respectively, while L_*ω*_ was calculated at 3 years (hereafter ‘L_3_’). The rVBGF model was fitted using the age (years) and length data (mm SL) of each sample and by constraining the curve to a length of settlement of 10 mm SL (Grandcourt et al. 2011). The best-fit model parameters, L_1_, L_3_ and L_6_, describing each dataset were determined by minimizing the negative log of the likelihood, given a probability density function with a Poisson distribution (Haddon 2001).

Mean maximum age (hereafter ‘Age_max_’) and mean maximum body size (hereafter ‘L_max_’) were estimated for both species in each region. Age_max_ and L_max_ were taken, respectively, as the average age (years) and body size (SL) of the 25% oldest or largest adult (i.e. ≥ 3 years old) individuals, respectively (Trip et al. 2008, 2014). Differences in mean Age_max_ and L_max_ between populations were analysed using linear models.

### 2.5 Individual annual growth estimation

To determine whether yearly growth rate differs within species between regions, annual increment widths were measured and compared between individuals. A subset of the total samples (148 and 144 otoliths of *L. ehrenbergii* and *P. maculosus*, respectively; Table 2) that showed clear edges between annual increments and with at least one completed annual increment (i.e. individuals ≥ 2 years old) were photographed and examined. As otolith growth is an appropriate representation of fish somatic growth in our species (i.e. significant positive relationship between otolith radius and individual body length, see Fig. S1a, b), increment widths were measured (mm) along the dorsal anti-sulcus axis (from core to edge) using MorphoJ (v1.06). Distances between the outer edges of each annual increment were taken to indicate the width of each consecutive growth increment (Fig. 2a, b). Through back calculation from age-at-capture (hereafter ‘AAC’) and year of capture, age (‘Age’) and calendar growth year (‘Year’) were respectively assigned to each increment (Table 3). The first two annual increments were not used in the analysis due to poor visualization in the inner region of the otolith, while the last (marginal) increments were excluded because they did not represent a full year of growth (Doubleday 2015; Martino et al. 2019). For each region, the sample size was at least five increment measurements per calendar year. In total, increment widths were measured from 148 *L. ehrenbergii* (Arabian Gulf = 58; Oman Sea =90) and 144 *P. maculosus* (Arabian Gulf = 107; Oman Sea = 37) individuals (Table 2). Increment widths encapsulated growth of individuals between 2002 and 2015 in *L. ehrenbergii* and between 1993 and 2016 in *P. maculosus* (Table 2).

**Table 2.**
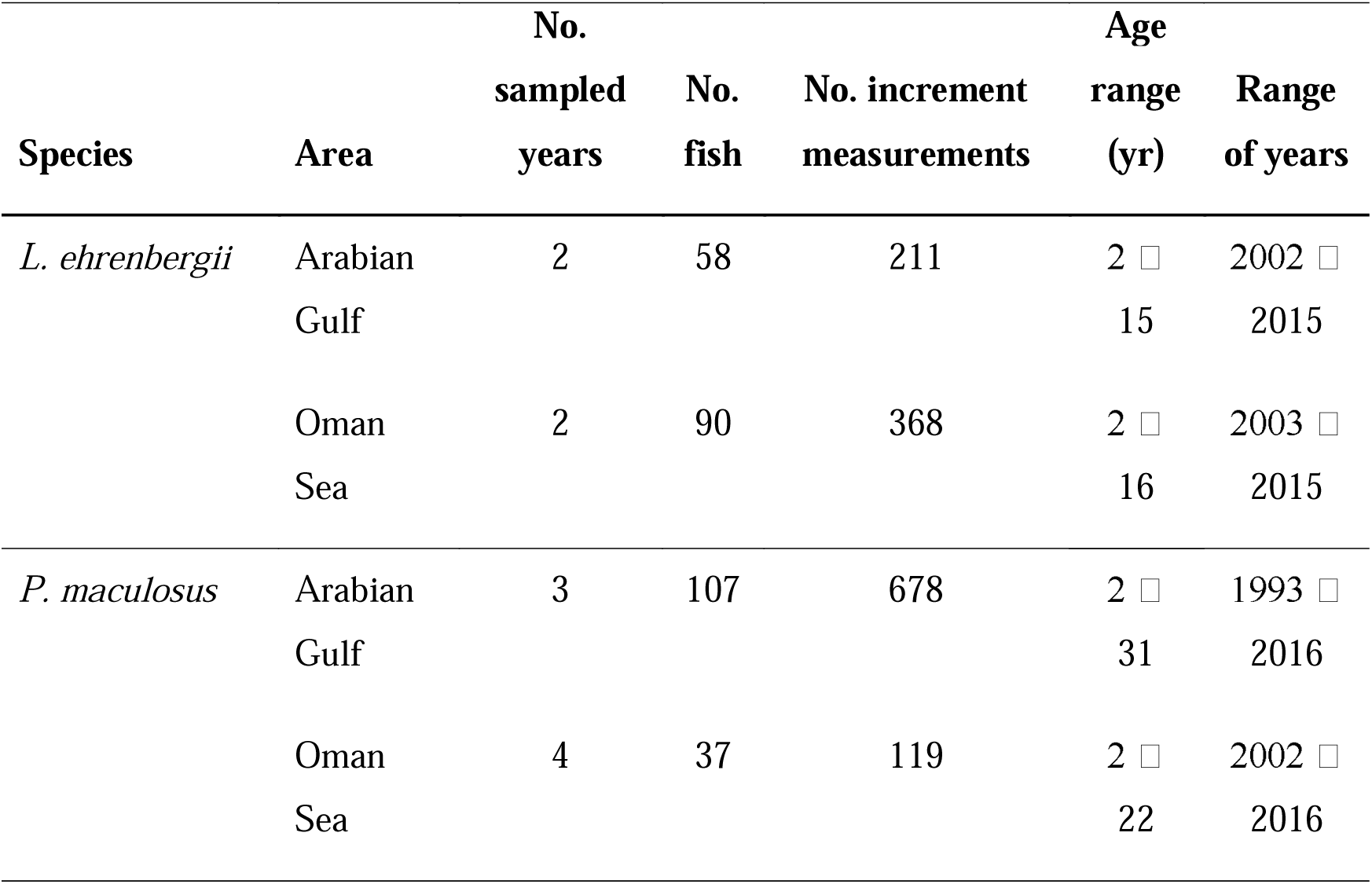
Summary of *L. ehrenbergii* and *P. maculosus* otolith samples used to develop sclerochronologies.

**Table 3.** Descriptions of predictor variables used in the mixed-effects modelling of *Lutjanus ehrenbergii* and *Pomacanthus maculosus*’s growth.

| Parameter | Description | Data range |
| --- | --- | --- |
| **Fixed intrinsic effects** |  |  |
| Age | Age in years when otolith annual increment was formed. |  |
| Year | Year in which the annual increment was formed. |  |
| AAC | Age in years at time of capture. Measure of potential selectivity on growth. |  |
| **Fixed extrinsic effects** |  |  |
| Mean SST | Mean annual sea surface temperature (°C). | 1993 □ 2016 |
| Max SST | Mean maximum sea surface temperature during the hottest month (Aug) (°C). | 1993 □ 2016 |
| Min SST | Mean minimum sea surface temperature during the coldest month (Feb) (°C). | 1993 □ 2016 |
| Salinity | Mean annual salinity (psu). | 1993 □ 2016 |
| Chl-a | Mean annual chlorophyll-a (mg m<sup>-3</sup>). | 2003 □ 2016 |
| **Random effects** |  |  |
| 1\|FishID | Unique identifier code for each fish. |  |
| 1\|Year | Calendar year in which the increment was formed, quantifies inter-annual growth variability. |  |
| Age\|FishID | Random age slope for FishID random intercept, allows for individual-specific differences in the growth–age relationship |  |
| Age\|Year | Random age slope for Year random intercept, allows for year-dependent differences in age-specific growth. |  |

### 2.6 Growth mixed modelling

Sources of annual growth variation in *P. maculosus* and *L. ehrenbergii* were investigated using a series of hierarchical mixed-effects models predicting variation in otolith increment widths (following Morrongiello and Thresher 2015). Growth was modelled separately for the two regions as to avoid the co-variation of multiple environmental factors with regions. All mixed modelling was performed in R (R core development team 3.5.1, 2019), using the packages lme4 (Bates et al. 2015), effects (Fox 2003), and AICcmodavg (Mazerolle 2019). Modelling could not be done for *P. maculosus* in the Oman Sea due to insufficient sample sizes.

To explore potential drivers of inter-annual growth variation within the model, we used both intrinsic and extrinsic predictors. Intrinsic predictors (i.e. intra-population drivers) were the fixed effects of Age and AAC, which account for expected age-related trends, bias, and selectivity (Morrongiello et al. 2012), while the random effects of FishID and Year were utilized (Table 3 and Supplementary materials – ‘intrinsic predictors’ for details). Extrinsic predictors (i.e. environmental predictors) were mean, maximum and minimum annual SST (°C) (‘mean-, max- and min SST’, respectively), annual mean salinity (‘salinity’, psu) and annual mean chlorophyll-a (‘chl-a’, mg m^-3^) (Table 3 and Supplementary materials – ‘extrinsic predictors’ for details). Due to the lack of sex data, sex was not used as growth predictor, nevertheless patterns of sexual dimorphism are known for our species (i.e. female individuals grow larger in *L. ehrenbergii*, while males grow larger in *P. maculosus*) (Grandcourt et al. 2010, 2011), and were taken into account during the interpretation of the results.

#### 2.6.1 Model selection

Growth, Age, and AAC were log-transformed to meet model assumptions. All variables (intrinsic and extrinsic) were mean centred to facilitate the convergence of the model and the interpretation of the random slopes (Morrongiello and Thresher 2015). Analyses were run with a two-stage approach: in the first stage, we built a base set of linear mixed models that encompassed a series of random effect and fixed intrinsic effect structures in order to identify the optimal intrinsic model (Tables S3 – S6 and Supplementary materials – ‘models selection’ for details of models selection, ranking and fitting). In the second stage of the analysis, we investigated whether environmental factors influenced growth. We extended the optimal intrinsic model for each region to include single or combinations between two environmental variables, which were fitted separately and ranked using Akaike’s information criterion corrected for small sample sizes (AIC_C_) (Burnham and Anderson 2004) and ΔAIC_c_ values (Table S7, S8). Interactions between SST effects and between SST and salinity were not included in the model selection as a high level of correlation was expected. Additionally, collinearity between all the remaining combinations was explored using variance inflation factor (VIF) and models with VIF above 3 were removed from comparison (Hair et al. 1998). Finally, linear temporal growth trends were investigated by adding Year as a fixed effect to the optimal intrinsic model.

## Results

### 3.1 Population-level growth, body size and life span

The age range of *L. ehrenbergii* individuals was similar between regions, ranging between 38 days to 15 years in the Arabian Gulf and 27 days to 16 years in the Oman Sea; whereas *P. maculosus* individuals’ age range differed between regions comprising of both younger and older individuals in the Arabian Gulf compared to the Oman Sea (9 days to 31 years and 81 days to 22 years, respectively) (Table 1).

Both *L. ehrenbergii* and *P. maculosus* had consistently smaller asymptotic size and maximum length (L_max_) in the Arabian Gulf compared with the Oman Sea (Fig. 3a, b; Table 1). Specifically, L_max_ of *L. ehrenbergii* and *P. maculosus* individuals was 19% (linear regression, F = 153.3, *df* = 1,49, *p* < 0.001) and 13% smaller (F = 9.968, *df* = 1,34, *p* = 0.003), respectively. Maximum age (Age_max_) was significantly lower in the Arabian Gulf compared to the Oman Sea for *L. ehrenbergii* (F = 4.362, *df* = 1,49, *p* = 0.042), but did not significantly differ between *P. maculosus* populations (F = 1.859, *df* = 1,34, *p* = 0.182) (Table 1).

**Fig. 3.3.**
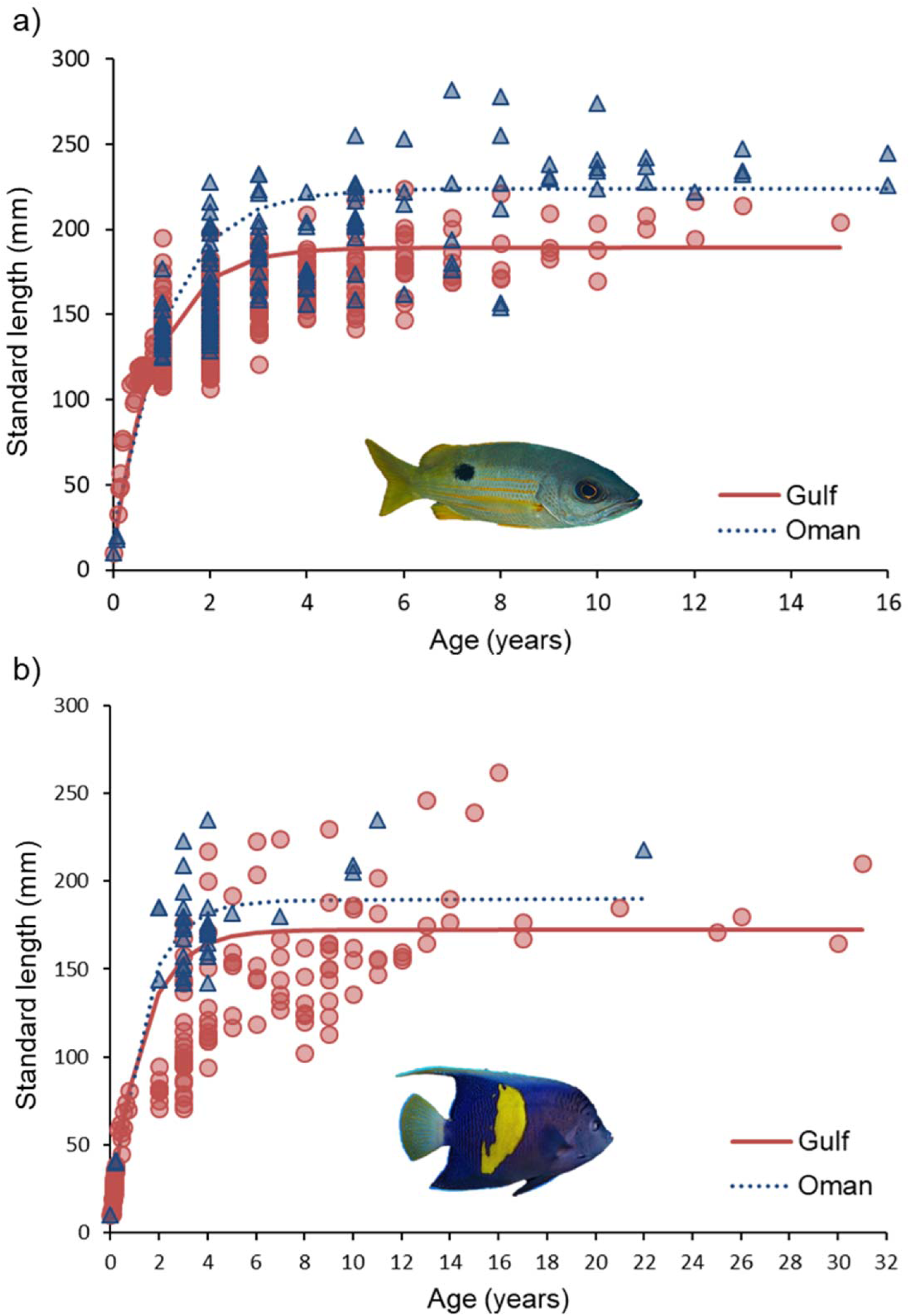
Comparison of re-parameterised version of the von Bertalanffy growth function (rVBGF) for *Lutjanus ehrenbergii* **(a)** and *Pomacanthus maculosus* **(b)** between the Arabian Gulf (‘Gulf’) and the Oman Sea (‘Oman’). Data points: red circles = Arabian Gulf, blue triangles = Oman Sea. Note different scales on the x-axis.

### 3.2 Sources of individual growth variation

#### 3.2.1. Extrinsic conditions

The magnitude and temporal trends of extrinsic parameters varied significantly between the Arabian Gulf and the Oman Sea (Fig. 4, Table S1). Both regions recorded relatively high inter-annual variation in min, mean, and max SST, while inter-annual variation in salinity was high in the Arabian Gulf but minimal in the Oman Sea. In addition, both regions recorded relatively high inter-annual variation in chlorophyll-a concentrations, which was stronger in the Oman Sea than the Arabian Gulf (Fig. 4a). Despite no difference in annual mean SST between regions, the Arabian Gulf had higher intra-annual SST variation, with temperature significantly colder in winter and warmer in summer compared to the Oman Sea, as well as higher annual mean salinities, and lower annual mean chlorophyll-a concentrations (Fig. 4b, Table S1).

**Fig. 4.**
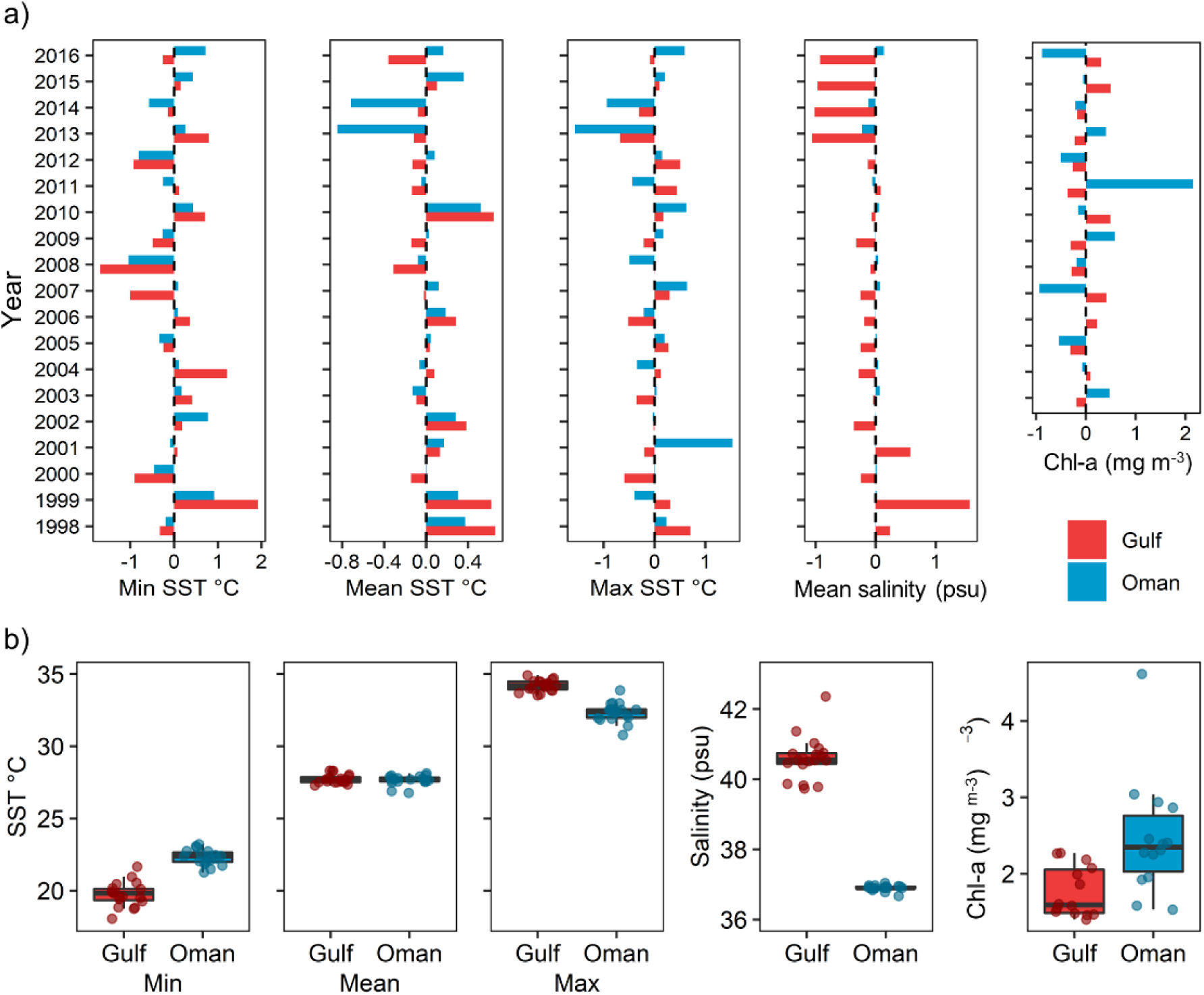
Intra- **(a)** and inter- **(b)** annual variability in sea surface temperature (min, mean and max), salinity and chlorophyll-a between the Arabian Gulf (‘Gulf’, red) and the Oman Sea (‘Oman’, blue). In **(a)** divergent bars quantify the positive/negative annual change of specific environmental variables during a given year from the individual mean (dashed vertical line) across all years. In **(b)** solid black line represents the median, with the box indicating the upper and lower quartiles, and whiskers representing the maximum or minimum observed value that is within 1.5 times the interquartile range of the upper or lower quartile, respectively. Dots are individual data points.

#### 3.2.2 Growth predictors

Across species and regions, growth declined with age and salinity (Table 4 and 5, see Table 4 for random effect structure details). Negative effects of salinity on growth rate were most apparent within the Oman Sea populations of *L. ehrenbergii* (−107.2% annual growth per unit increase in annual mean salinity [psu^-1^]), while Arabian Gulf populations of *L. ehrenbergii* and *P. maculosus* also showed lower growth with higher levels of salinity (−30.3% and - 5.1% annual growth per unit increase in annual mean salinity [psu^-1^], respectively) (Fig. 5a and Table 5).

**Table 4.** Optimal model structures derived by ranking a series of increasingly complex mixed-effects models.

| Species | Region | Intrinsic model | Fixed effects | Random effects | Best extrinsic covariate structure |
| --- | --- | --- | --- | --- | --- |
| *L. ehrenbergii* | Arabian Gulf | 2a2 | Age + AAC | Age\|FishID + 1\|Year | Salinity + max SST |
|  | Oman Sea | 2a2 | Age + AAC | Age\|FishID + 1\|Year | Salinity |
| *P. maculosus* | Arabian Gulf | 3a1 | Age | Age\|FishID + Age\|Year | Salinity + mean SST |
|  | Oman Sea | NA | NA | NA | NA |

In *Lutjanus ehrenbergii*, the best intrinsic model was similar across regions and included Age and AAC with a random Age slope for FishID. This indicated that the growth-age relationship differed among individuals, while the random intercept for Year indicated growth varied among years. In *Pomacanthus maculosus* in the Arabian Gulf the best intrinsic model included only Age as fixed intrinsic factor with random Age slope for FishID and random Age slope for Year, indicating the age-growth relationship differed among years.

**Table 5.**
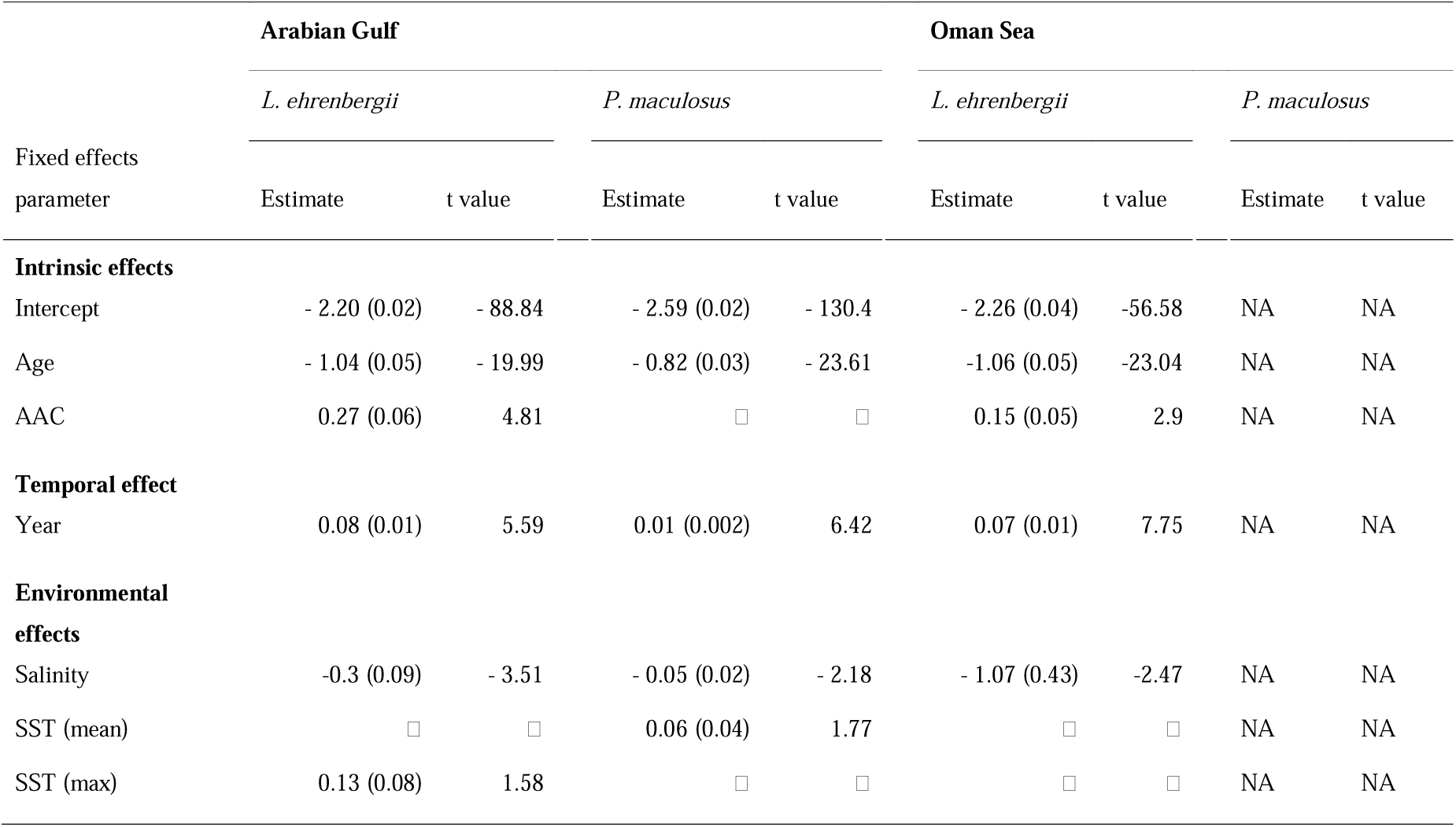
Fixed effect parameter estimates (± SE) and test statistics for optimal models describing intrinsic, temporal and environmental effects on *Lutjanus ehrenbergii* and *Pomacanthus maculosus*’ growth (see Tables S1 □ 8 for model selection and base models’ detail).

| Fixed effects<br>parameter | Arabian Gulf |  |  |  | Oman Sea |  |  |  |
| --- | --- | --- | --- | --- | --- | --- | --- | --- |
|  | *L. ehrenbergii* |  | *P. maculosus* |  | *L. ehrenbergii* |  | *P. maculosus* |  |
|  | Estimate | t value | Estimate | t value | Estimate | t value | Estimate | t value |
| **Intrinsic effects** |  |  |  |  |  |  |  |  |
| Intercept | - 2.20 (0.02) | - 88.84 | - 2.59 (0.02) | - 130.4 | - 2.26 (0.04) | -56.58 | NA | NA |
| Age | - 1.04 (0.05) | - 19.99 | - 0.82 (0.03) | - 23.61 | -1.06 (0.05) | -23.04 | NA | NA |
| AAC | 0.27 (0.06) | 4.81 |  |  | 0.15 (0.05) | 2.9 | NA | NA |
| **Temporal effect** |  |  |  |  |  |  |  |  |
| Year | 0.08 (0.01) | 5.59 | 0.01 (0.002) | 6.42 | 0.07 (0.01) | 7.75 | NA | NA |
| **Environmental effects** |  |  |  |  |  |  |  |  |
| Salinity | -0.3 (0.09) | - 3.51 | - 0.05 (0.02) | - 2.18 | - 1.07 (0.43) | -2.47 | NA | NA |
| SST (mean) |  |  | 0.06 (0.04) | 1.77 |  |  | NA | NA |
| SST (max) | 0.13 (0.08) | 1.58 |  |  |  |  | NA | NA |

There was little consistent effect of increasing temperature on growth rate between regions, though regional patterns were apparent (Fig. 5b and Table 5). Water temperature within the Arabian Gulf was positively correlated to growth in both species, with increases in max SST having a predicted effect on *L. ehrenbergii*’s growth rate of + 12.7% per °C, and increases in mean SST an effect of + 6.5% per °C in *P. maculosus*. There was no effect of temperature on the Oman Sea populations of *L. ehrenbergii*.

**Fig. 3.6.**
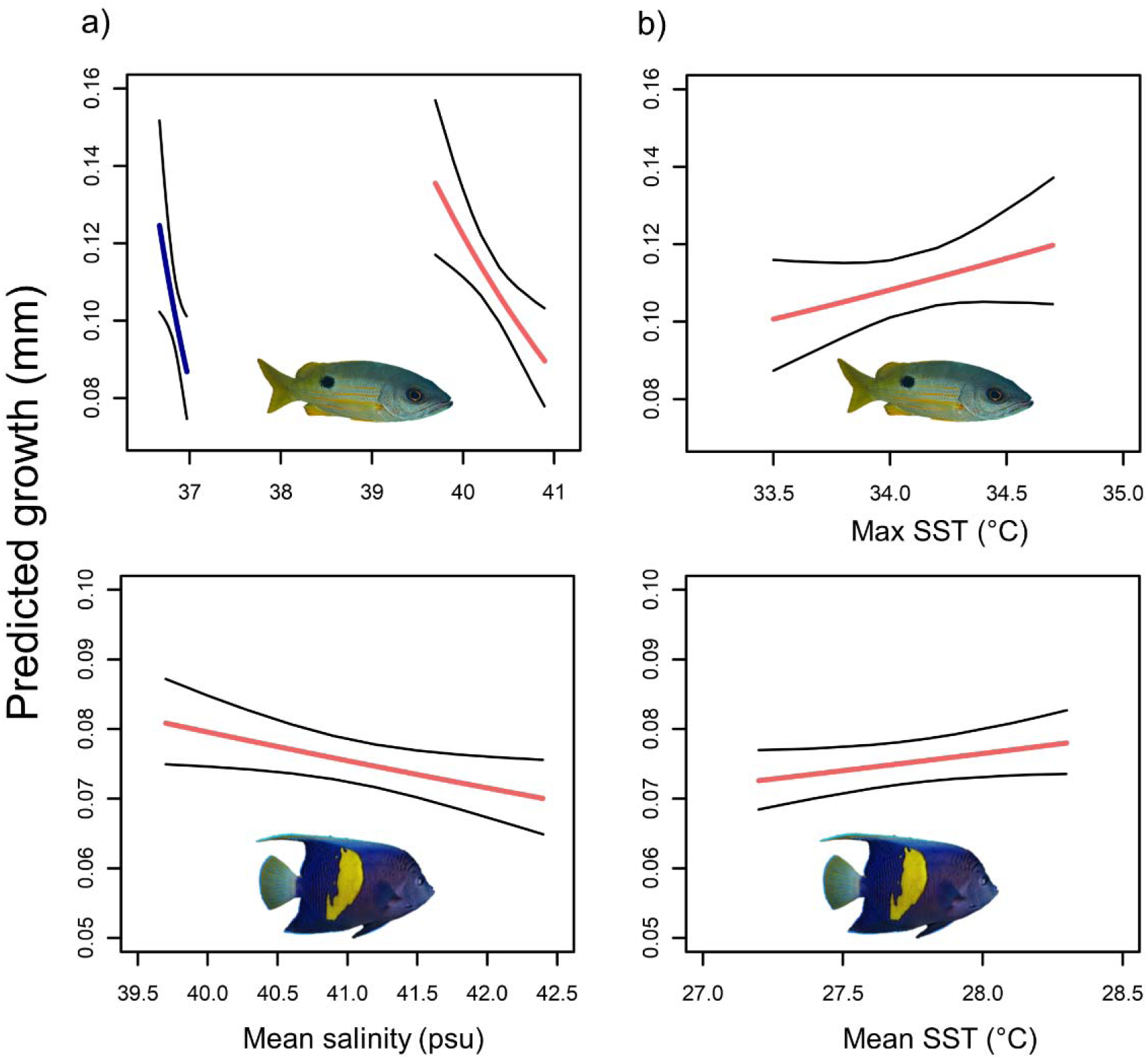
Predicted effect of annual mean salinity **(a)** and max and mean sea surface temperature (SST) **(b)** on the otolith incremental growth of *Lutjanus ehrenbergii* and *Pomacanthus maculosus* populations between the Arabian Gulf (red lines) and Gulf of Oman (blue lines). Black lines indicate 95 % CI. Note different scales on the y and x-axis. Predicted effects refer to the model outputs described in Table 3.4 and 3.5.

## Discussion

We compared the growth of two common coral reef fish *L. ehrenbergii* and *P. maculosus* between the environmentally extreme Arabian Gulf and comparably benign Oman Sea, and examined the impact of temperature, salinity and primary productivity on the somatic growth of populations within each region. We found that both species were significantly smaller at age and attained a smaller maximum body size within the Arabian Gulf than within the Oman Sea. However, contrary to the expectation that extremes in temperature would be the main driver of somatic size reduction (i.e. consistent with the TSR), reductions in body size were mainly related to variation in salinity. Such results indicate that salinity may be a vital determinant of both species’ growth trajectories. In comparison, temperature had a slightly positive effect on growth rates of Arabian Gulf populations of both species, with no measurable effect on Oman Sea fish populations.

In our study, growth declined with higher mean annual salinity across species and regions, indicating the impacts on demography of a highly saline environment are a reduction and, ultimately, truncation of life history. Indeed, together with temperature, food availability and photoperiod, salinity is known to be a major factor in determining fish development and growth (Boeuf and Payan 2001). Moreover, as osmoregulation is energetically costly (i.e. between 2–30% of daily energy expenditure in marine fish) (Boeuf and Payan 2001; Ern et al. 2014; Kultz 2015), changes in salinity may result in significant energy diversion from growth (Boeuf and Payan 2001). Here, we did not observe the expected crossing of growth curve trajectories between populations living in different thermal regime as predicted by the TSR (i.e. faster juvenile growth but earlier asymptotic growth for fish living in warmer environments) (Trip et al 2014), suggesting that salinity and perhaps even productivity are exhibiting a strong influence on differences in growth between regions. Interestingly, growth rates of populations within the Oman Sea, where salinity conditions are predominantly stable, were impacted more strongly by salinity fluctuations compared to populations within the Arabian Gulf, suggesting Arabian Gulf’s populations may have higher capability of acclimation to seasonal and interannual changes in environmental conditions (Rummer and Munday 2017; D’Agostino et al. 2019). While numerous studies have examined the implications of increasing temperature and temperature variability on coral reef fish growth (e.g. Munday et al. 2008; Messmer et al. 2017; Taylor et al. 2019), to our knowledge there has been no investigation of the combined effect of increasing temperature and salinity on coral reef fish demography. This is despite the evidence of synergism between the two stressors (Claireaux and Lagarde 1999; Jian et al. 2003) and the likely occurrence of both stressors in already dry regions and hyper-saline (*sensu* saltier than ocean salinity) semi-enclosed seas in the near future (such as the Arabian Gulf, Red Sea and Mediterranean Sea) (Durack et al. 2012; Skliris et al. 2014; Zika et al. 2018).

Positive effects of high SST on growth rates of both species within the Arabian Gulf may show that high summer water temperatures do not exceed the population’s typical temperature range in this region. For example, there was a positive effect of increased maximum SST on growth rate of *L. ehrenbergii*, potentially indicating that the Arabian Gulf’s summer temperature of > 34 °C may not exceed the thermal optimum for this species. Indeed, *L. ehrenbergii* abundance and predatory activity appear to be highest in summer (D’Agostino et al. 2019), suggesting that *L. ehrenbergii* may still have the aerobic capacity to perform ecological tasks (e.g. swimming, feeding) during the Arabian Gulf’s extreme summer temperature. Additionally, the positive effect of mean SST on Arabian Gulf populations of *P. maculosus* may suggest that intermediate temperatures experienced during milder winters or slightly warmer springs or autumns may benefit the growth rate of this species (Johansen et al. 2015; Djurichkovic et al. 2019).

The smaller body sizes observed in Arabian Gulf fishes of both species may represent a life-history trade-off between metabolic demands (i.e. increased osmoregulatory cost) and size, with likely flow-on effects to population structure. Although not universally accepted, the gill-oxygen limitation theory (GOLT) states that as gills function as a two-dimensional surface with growth limited by geometrical constraints, any three-dimensional increases in fish body size, and consequent increases in oxygen demand, may not be met by adequate oxygen supply (Pauly 1981; Pauly and Cheung 2018b, but see Audzijonyte et al. 2019). Consequently, larger-bodied individuals may be unable to compensate for increased metabolic demands associated with high salinity and temperature, due to the incapability of the respiratory system to supply enough oxygen. While we did not observe a strong negative effect of max SST (when dissolved oxygen is at its lowest and the oxygen demand at its highest) on fish growth (Shapiro Goldberg et al. 2019), large-bodied individuals are expected to have limited capacity to increase mass-specific maximum metabolic rate in warmer conditions (Messmer et al. 2017), and to be more likely to approach their maximum physiological capacity (Pauly and Cheung 2018a). Hence, reduced individual body size in the Arabian Gulf may represent a life-history trade-off, whereby survival is enhanced through smaller body size and, therefore, reduced metabolic demands.

Although reductions in body size may play an important role in coping with variable and extreme temperature and salinity, other mechanisms may be involved in facilitating population stability. For example, recent work has highlighted the importance of behavioural and feeding plasticity in coping with the Arabian Gulf’s extreme environmental conditions. The pale-tail damselfish (*Pomacentrus trichrourus*) appears able to mitigate bioenergetic inefficiency associated with the fluctuating and extreme water temperature within the Arabian Gulf by downregulating costly activities during winter and summer, while upregulating activity and increasing energy stores in spring (D’Agostino et al. 2019). In this respect, *P. trichrourus, P. maculosus* and *Pomacentrus aquilis* all show a degree of feeding plasticity in the Arabian Gulf (Shraim et al. 2017; D’Agostino et al. 2019), suggesting that such plasticity in feeding may be an important factor in understanding growth rate and overall body size of all three species.

Reductions in fish body size, within the magnitude of lowered size reported in the present work, are expected to have substantial consequences for trophic interactions, ecosystem function, fisheries and global protein supply (Shackell et al 2010; Cheung et al. 2013). Fish body sizes are already reducing due to intensive fishing pressure (Stergiou 2002), while there is also evidence to show the potential additive effect of climate change and oxygen limitation on individual (and therefore population) body size spectrum (Cheung et al. 2013; van Rijn et al 2017). Notably, theoretical models predict that small reductions in individual body size (i.e. 4% over 50 years) may lead to a 50% increase in mortality as well as a 5 □ 35% reduction in biomass and catch (Audzijonyte 2013). However, most importantly, as fish fecundity is largely correlated with body size, lowered body size is expected to have substantial negative consequences on population reproductive output, replenishment and long-term persistence (Baudron 2014), which will have cascading effects on the wider ecosystem, potentially impacting coastal economies and threatening food security.

Understanding the mechanisms by which fishes endure the Arabian Gulf’s extreme and variable environment will be vital to understand how low latitude coral reef fish populations may cope with predicted changes in their environment. The present work shows that sub-tropical fish communities can persist within extreme environmental conditions, albeit with substantial trade-offs to their life history and demographic structure; individuals of both *L. ehrenbergii* and *P. maculosus* showed smaller size-at-age and lower maximum size in the Arabian Gulf compared with conspecifics in the Oman Sea. Our results suggest that, in order to predict the effect of climate change on fish demography, the combined effect of osmoregulatory and thermal stress needs to be considered, especially in regions with limited oceanic exchange and predicted increase SST and evaporation. Ultimately, to tease apart the effect of extreme temperature and salinity on growth of Arabian Gulf fish populations and establish the role of other potential coping mechanisms (i.e. feeding and behavioural plasticity) in mitigating environmental stressors, laboratory-based physiological-behavioural experiments combining multiple stressors and conditions, while assessing detailed energy budget, are needed.

## Acknowledgements

This work was funded by a PhD scholarship from the University of Nottingham to D. D. and supported by NYU Abu Dhabi Marine Biology Core Technology Platform. Field fish collection was carried with permission of the Environment Agency, Abu Dhabi (protocol No. EAD-TMBS-RP-0) and according to the NYU-Abu Dhabi animal ethics guidelines. We also thank A. D. C. MacColl and the anonymous reviewers for useful advice and constructive suggestions. Corresponding author’s ORCID ID: https://orcid.org/0000-0003-2291-5749.

